# Enhanced donor antigen presentation by B-cells predicts acute cellular rejection and late outcomes after transplantation

**DOI:** 10.1101/2022.10.17.511155

**Authors:** Chethan Ashokkumar, Mylarappa Ningappa, Vikram Raghu, George Mazariegos, Brandon W. Higgs, Paul Morgan, Lisa Remaley, Tamara Fazzolare, Pamela Holzer, Kevin Trostle, Qingyong Xu, Adriana Zeevi, James Squires, Kyle Soltys, Simon Horslen, Ajai Khanna, Armando Ganoza, Rakesh Sindhi

**Author notes:** **Corresponding Author** Rakesh Sindhi, MD, FACS, Professor of Surgery, University of Pittsburgh, UPMC Children’s Hospital of Pittsburgh, 4401 Penn Avenue, FP-6/Transplant/Rm 6140, Pittsburgh, PA 15224 /6117.

## Abstract

**Purpose:** Enhanced B-cell presentation of donor alloantigen relative to presentation of HLA-mismatched reference alloantigen is associated with acute cellular rejection (ACR), when expressed as a ratio called the antigen presenting index (API) in an exploratory cohort of liver and intestine transplant (LT, IT) recipients.

**Methods:** To test clinical performance, we measured the API using the previously described 6-hour assay in 84 LT and 54 IT with median age 3.3 years (0.05-23.96). Recipients experiencing ACR within 60 days after testing were termed rejectors.

**Results:** We first confirmed that B-cell uptake and presentation of alloantigen induced and thus reflected the alloresponse of T-helper cells, which were incubated without and with cytochalasin and primaquine to inhibit antigen uptake and presentation, respectively. Transplant recipients included 76 males and 62 females. Rejectors were tested at median 3.6 days before diagnosis. The API was higher among rejectors compared with non-rejectors (2.2 ± 0.2 vs 0.6 ± 0.04, p-value=1.7E-09). In logistic regression and ROC analysis, API ≥ 1.1 achieved sensitivity, specificity, positive and negative predictive values for predicting ACR in 99 training set samples. Corresponding metrics ranged from 80-88% in 32 independent post-transplant samples, and 73-100% in 20 independent pre-transplant samples. In time-to-event analysis, API ≥ 1,1 predicted higher incidence of late DSA after API measurements in LT (p=0.011) and graft loss in IT recipients (p=0.008), compared with recipients with API<1.1, respectively.

**Conclusion:** Enhanced donor antigen presentation by circulating B-cells predicts rejection after liver or intestine transplantation as well as higher incidence of DSA and graft loss late after transplantation

## Introduction

The presentation of donor antigen by B-cells presents novel opportunities to further characterize, predict and treat cellular and humoral rejection in transplant recipients. B-cells present antigen to T-helper cells (Th), resulting in Th-mediated alloresponses. Th-mediated help is essential for B-cell activation and maturation to antibody secreting plasma cells, and for activation and memory formation in T-cytotoxic cells (Tc) (1-3). Activated B-cells can also recruit other B-cells to produce antibodies (4, 5). The CD40-CD154 interaction facilitates this communication within and between cell compartments, and depends in part on the induction of CD40 on Tc and Th (6, 7). Reflecting the clinical import of these interactions, donor-antigen-specific B-cells, which express CD154 in lymphocyte co-culture are enhanced during acute cellular rejection after intestine transplantation, and are strongly correlated with allospecific CD154+Tc-memory cells (TcM), and with donor-specific anti-HLA antibodies (DSA) (8). B-cell presentation of alloantigen may be central to these observations. In mice with major histocompatibility complex (MHC) class II-deficient or *HLA-DM*-deficient B-cells, cardiac allograft survival was enhanced, and was accompanied by decreased T-cell activation and alloantibody production (9). HLA-DM facilitates the exchange of peptides loaded on MHC II during antigen presentation by B-cells (10). Interestingly, peptides which complex more stably with MHC II are more likely to elicit Th activation, a pre-requisite for subsequent cellular and humoral alloresponses (10).

We have previously described a novel assay which measures donor-antigen uptake by B-cells, and identifies rejection-prone children with liver transplantation within a few hours (11). This test system measures uptake of donor antigenic lysate in lieu of antigen presentation and is based on previous studies by others which show a direct correlation between uptake of non-allogeneic antigens and T-cell activation, the downstream effect of antigen presentation (12-15). We have used this assay to evaluate the role of the *HLA-DOA* gene, which inhibits HLA-DM-mediated peptide exchange on MHC II molecules during antigen presentation by B-cells (16). In genetic association studies, single nucleotide variants in the regulatory regions of *HLA-DOA* were associated with liver transplant rejection in children. The *HLA-DOA* gene inhibits antigen presentation in B-cells (17).

Here, we evaluate whether B-cell uptake of alloantigen represents presentation of alloantigen and the downstream T-cell response. Further, we establish the performance of this test system as a predictor of rejection and late DSA in a larger cohort of liver and non-liver allograft recipients. The appearance of DSA late after transplantation is associated with graft loss (18, 19).

## Methods

*Human subjects*: Ficoll-purified PBL were extracted from blood samples from three healthy adult human subjects and a total of 138 liver or intestine transplant recipients. All subjects were sampled after IRB-approved informed consent, and purified PBL archived in liquid nitrogen. Archived samples for this study were selected based on availability, and proximity to a defined biopsy-proven event or a clinical visit for surveillance within a 60-day period after sampling.

### B-cell antigen uptake, presentation and the T-cell alloresponse

To understand whether an antigen uptake assay was reflective of antigen presentation, we first modeled indirect antigen presentation by incubating antigenic lysate with an HLA-non-identical responder PBL mixture consisting of purified B-cell and Th. For this experiment, a) fluorochrome-labeled antigenic lysate labeled with carboxyfluoresciensuccinimidylester (CFSE) was prepared from peripheral blood lymphocytes (PBL), b) responder B-cells and Th were purified individually using Miltenyi magnetic beads, and reconstituted to make up responder PBL. Three types of responder B-cells were used in each of three replicates of this experiment: untreated B-cells, B-cells pre-treated with cytochalasin D, an inhibitor of antigen uptake, and B-cells pre-treated with primaquine, an inhibitor of antigen presentation. The frequency of CD19+B-cells which express fluorochrome-labeled lysate, and CD4+Th which express CD154 were measured by flow cytometry.

### Predicting rejection in transplant recipients

PBL from children with liver or intestine transplantation were tested with our previously described antigen presenting assay in which the uptake of donor and HLA-non-identical third-party (reference) antigen was measured in parallel reaction conditions, as described (11). The results were expressed as a ratio called the antigen presenting index (API).

### Statistical methods

In 99 training set samples from 99 subjects with liver or intestine transplantation, exhaustive forward, backward, and stepwise logistic regression (LR) and receiver-operating-characteristic (ROC) curve analysis was used to identify a threshold API associated with rejection within the 60-day period after the assay, as well as sensitivity, specificity, positive and negative predictive values, as described previously (20-22). Covariates in the model included gender: male and female, race: white and non-white, induction: no induction/ induction, tacrolimus whole blood concentration (FKWB). Age at transplantation was included as a continuous variable. The performance of this threshold API was evaluated in independent post-transplant and pre-transplant samples. We used SPSS version 27 (IBM Corporation, NY) to perform descriptive statistics, between-group comparisons with t-tests, and LR-ROC analyses. We used Stata v17 (StataCorp, College Station, TX) to perform survival analysis. We built a Cox-proportional hazards stratified on organ transplanted to determine the association between API and DSA development.

## Results

### Human subjects

General demographics for 138 subjects are shown in Table S1. These subjects included 84 liver transplant recipients including one who also required renal transplantation for congenital hepatic fibrosis and polycystic kidney disease, 29 recipients of a liver-inclusive intestine allograft, and 25 recipients of an intestine allograft of whom one also received a pancreas allograft.

#### The indications for liver transplant

were alpha-1-antitrypsin deficiency-2, autoimmune hepatitis-1, Alagille’s syndrome-4, acute liver failure-4, autoimmune hepatitis-1, Biliary atresia-20, Caroli’s disease-1, cryoptogenic cirrhosis-3, Criggler-Najjar syndrome-3, cystic fibrosis-1, congenital hepatic fibrosis-1, familial cholestasis-2, glycogen storage disease-1, Malignancy-8 (including hepatoblastoma-5, embryonal sarcoma-2, malignant rhabdoid-1), MSUD-12, Neonatal hepatitis-1, primary hyperoxaluria-1, primary sclerosing cholangitis-1, tyrosinemia-1, urea cycle defect-6, Wilson’s disease-3

#### The indications for intestine transplant

without or with liver or pancreas allograft were gastroschisis-9, Hirschsprung’s disease-5, trauma-2, jejunal atresia-3, necrotizing enterocolitis-6, pseudo-obstruction-6, unknown 3, volvulus-14, and secretory diarrhea group of diseases-6 (including microvillous inclusion disease, tufting enteropathy).

#### Outcome groups

Rejectors were sampled at a mean interval of 3.6 (range 0 to 30) days before diagnosis and treatment of rejection. Rejection diagnosed with Banff criteria was present in all but one subject. This single training-set subject was diagnosed as having clinical rejection, based on elevated liver function tests, absence of bile duct obstruction and arterial thrombosis on ultrasound, and normalization of liver function tests with steroid treatment.

#### Subject cohorts for performance testing

Ninety-nine post-transplant samples from 99 recipients made up the training set. Model predictions developed in this training set were validated in 32 post-transplant samples from 32 independent subjects, and 20 pre-transplant samples. These 20 single pre-transplant samples were obtained from 11 subjects in the training set, two in the validation set, and 7 independent subjects. Subject characteristics are shown in Table S1.

### B-cell alloantigen uptake reflects B-cell presentation of alloantigen and Th alloresponse

Uptake of HLA-non-identical CFSE-labeled antigenic lysate by B-cells was inhibited by pre-treatment of B-cells with cytochalasin D, an inhibitor of antigen uptake but not by primaquin, an inhibitor of antigen presentation in three replicates of this experiment. However, downstream Th alloresponse measured with CD154+Th cells was inhibited by B-cell pretreatment with either inhibitor. Bar diagrams in Figure 1a summarize results of three replicates of this experiment. Thus, antigen uptake measured by our assay is also representative of the final step of alloantigen presentation by B-cells, and the downstream Th alloresponse.

**Figure 1a:**
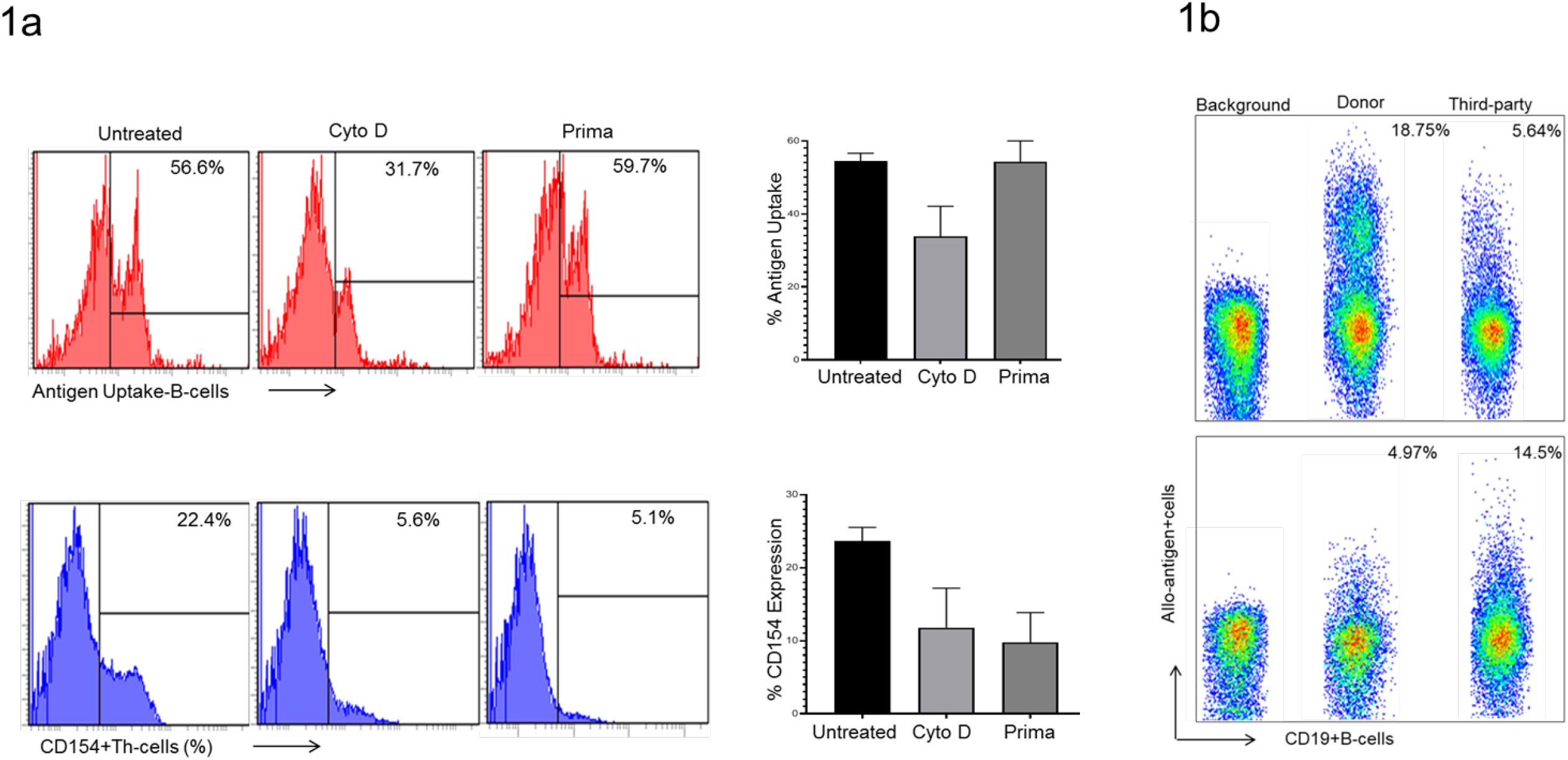
Alloantigen uptake, presentation and Th alloresponse. B-cells purified from normal human PBL were treated with Cytochalasin D (Cyto D) or Primaquine (Prima) and reconstituted with purified CD4 Th from the same subject into a responder cell mixture. Stimulation was performed with HLA-non-identical CFSE-labeled antigenic lysate. *Upper panel*. Histograms show that uptake of HLA-non-identical CFSE-labeled antigenic lysate by B-cells. *Lower panel*. Histograms show the downstream T-cell alloresponse measured as frequencies of CD154+CD4Th cells after each treatment. Bar diagrams on right summarize results from 3 replicates. **Figure 1b:** B-cell alloantigen presentation in a rejector and non-rejector. Three scatterplots each from a rejector (upper three plots) and a non-rejector (lower three plots) show frequencies of responder B-cells that present fluorochrome-labeled donor alloantigen (donor) and dye-labeled HLA-non-identical (reference) alloantigen. The background indicates recipient B cells cultured without either alloantigen.

### An API threshold of 1.1 or greater predicts acute cellular rejection in transplant recipients

In LR analysis, the optimal model identified API as a single variable predictor of rejection in the training set. ROC curve analysis identified an API of 1.06 as having the optimal balance of sensitivity and specificity for prediction of rejection. For corroboration, API measurements from the 54 liver transplant and 45 intestine transplant recipients in the training set were subjected to separate LR and ROC analyses, using the same covariates applied to the combined analyses. The optimal model once again identified API as the best predictor of rejection outcome at or above threshold values of 1.06 and 1.1085 respectively. We chose the threshold API of 1.1 at or above which rejection is predicted as a common threshold for all organs and applied it to 32 post-transplant and 20 pre-transplant validation samples to determine test performance.

*Test performance*: A higher proportion of B-cells presented donor antigen relative to reference alloantigen, leading to a significantly greater API among rejectors compared with non-rejectors (2.2±0.2 vs 0.6±0.40, p-value=1.7E-09, Figure 1b) in the combined cohort of 151 training and validation set samples from 138 liver and intestine transplant recipients. Logistic regression and ROC analysis showed that API ≥ 1.1 predicted ACR in the 99 training set samples with sensitivity, specificity, positive predictive value and negative predictive values and corresponding 95% confidence intervals (CI) of 93% (CI-76-99), 97% (CI-89-100), 93% (CI-76-99) and 97% (CI-88-99), respectively. In 32 independent post-transplant blinded validation set samples, respective performance metrics for the threshold API ≥ 1.1 were 86% (CI-57-98), 83% (CI-59-96), 80% (CI-52-96) and 88% (CI-64-99), respectively. In 20 pre-transplant samples respective performance metrics for the threshold API ≥ 1.1 were 73% (CI-39-94), 100% (CI-55-100), 100% (CI-52-100) and 75% (CI-43-95), respectively (Table 1). These results are summarized in Table 1 and Figure S1.

**Table 1.**
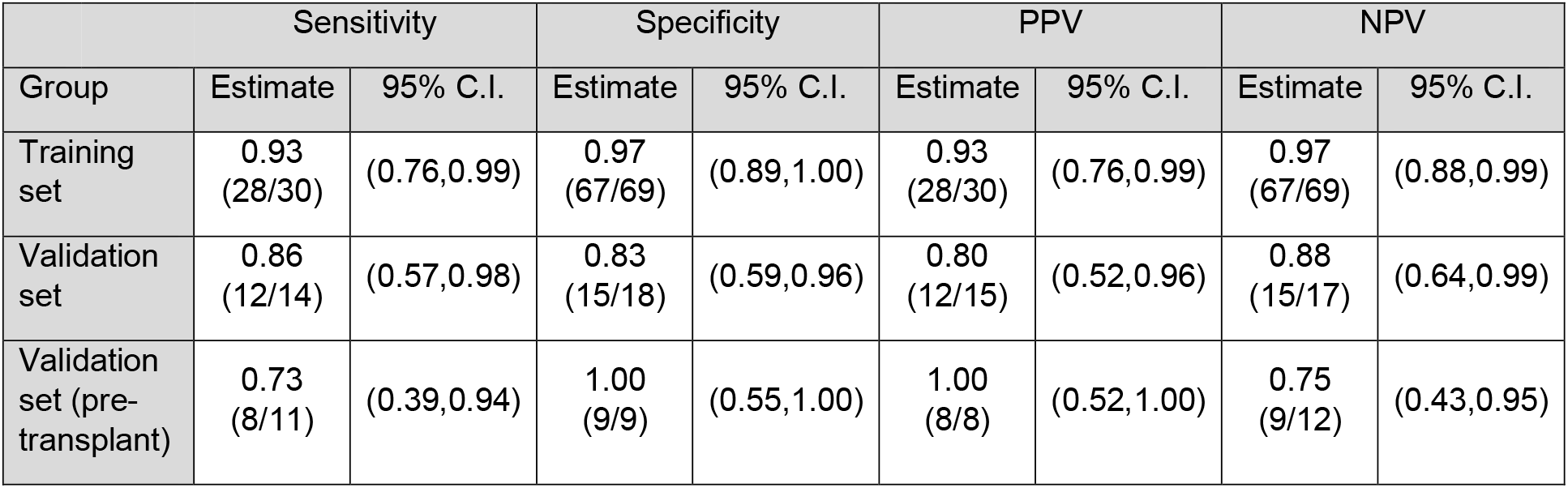
Summary of test performance measured in the various cohorts based on the API threshold of 1.1 or greater. Unless otherwise stated, post-transplant samples were used in the training set and validation set. PPV-positive predictive value, NPV-negative predictive value, CI-confidence interval

### Enhanced donor antigen presentation by B-cells precedes future development of DSA

DSA measurements were available for 85 of 138 subjects with post-transplant API measurements. These 85 recipients included 32 with an API ≥ 1.1 indicative of increased presentation of donor antigen, and 53 children with API < 1.1. B-cell API was measured at median 411 (5-5817) days after transplantation. In recipients with API ≥ 1.1, the incidence of DSA was higher at 20/32, 63% versus 17/53, 32%, p=0.006, Fisher’s exact, compared with those with API<1.1. Time to DSA in the respective groups was lower with median 2401 days (392-3539) versus 3122 days (825-8282), in recipients with high API compared with those with API < 1.1 (p < 0.01, K-M test). In the subcohort of 55 LT recipients, those with API ≥ 1.1 had a higher incidence, 19/28 (68%) versus 10/27 (37%), and earlier detection of DSA at 2989 days (1359-6991 versus 3204 days (825-8282) in those with API <1.1 (p=0.011, K-M test). Among IT recipients, DSA were present in 1/4 (25%) recipients with API > 1.1 and 6/26 (23%) recipients with API <1.1 (p=NS). In Kaplan-Meier analysis, greater risk of DSA development was seen in those with API > 1.1 among all recipients (p<0.01) and those receiving livers alone (p=0.011) (Figure 2 and Figure S2). Accounting for organ type, API > 1.1 was associated with 2.29 (95% CI 1.15 - 4.59, p=0.019) times the risk of DSA development over time.

**Figure 2:**
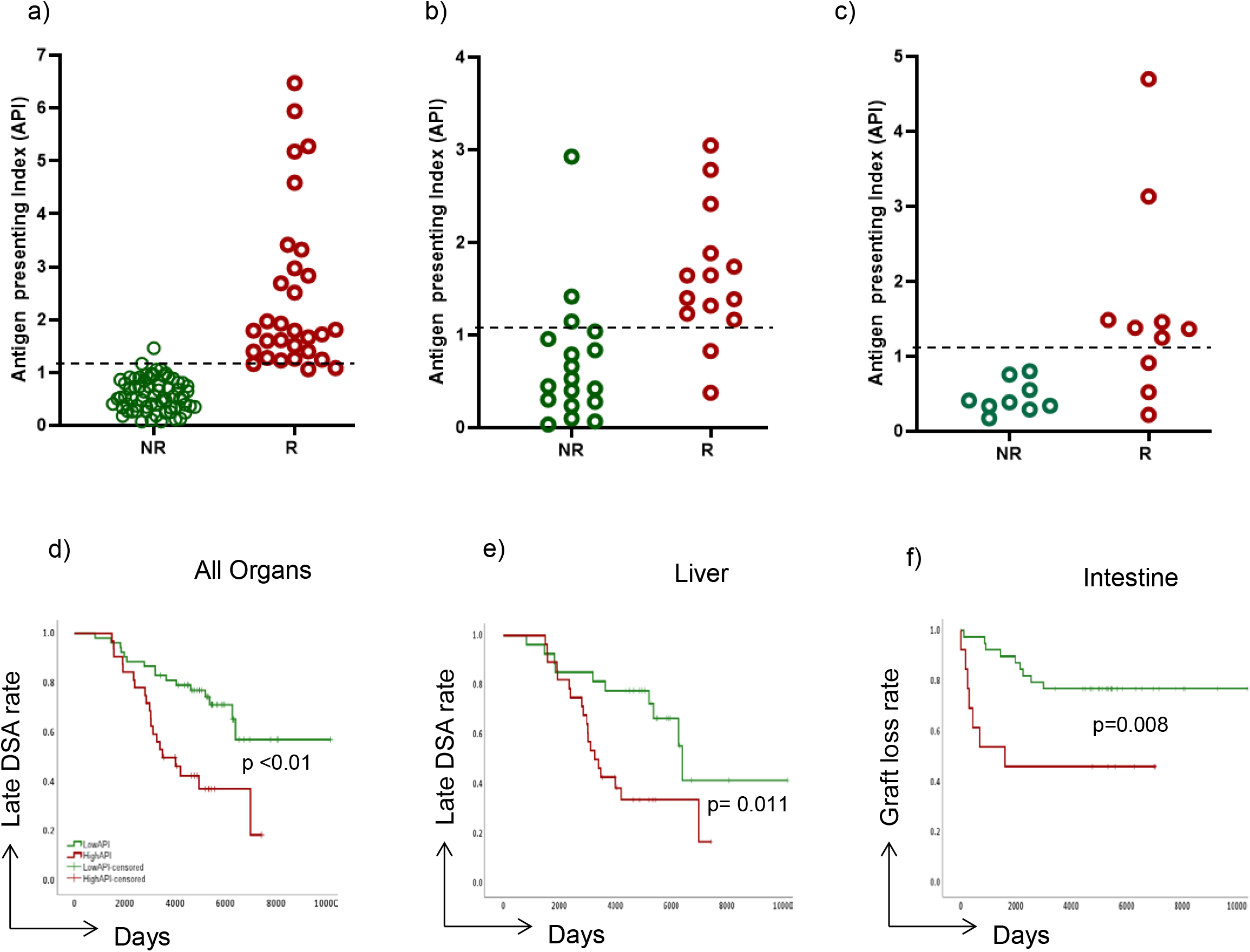
Test performance. *Upper row*. Dot plots show API values for rejectors and non-rejectors in a) all post-transplant training set samples, b) all post-transplant validation set samples and c) all pre-transplant validation set samples. *Lower row*. K-M plots show time to late donor-specific anti-HLA antibody (DSA) measurements d) 85 of 131 total study subjects for whom DSA measurements were available, and e) Fifty five LT recipients. f) K-M plot shows graft loss rate in 52 IT recipients. Each plot shows two subgroups, recipients with API <1.1 (green) and those with API ≥ 1.1 (red).

### Enhanced donor antigen presentation by B-cells is associated with Graft loss in Intestine transplant recipients

Of 131 LT and IT recipients who contributed training and validation sets of post-transplant samples, 20 experienced graft loss. These graft losses occurred in 9/45 recipients with API ≥ 1.1 and 10/86 with API < 1.1, at median (range) 1626 days (190-6250) versus 2390 days (228-4661), respectively (p=0.235, NS, K-M test). In Kaplan-Meier analysis, the risk of graft loss was greater for intestine recipients with API >1.1 (p<0.01). Graft losses occurred earlier at median 727 days (190-6250) in 7/13 (54%) recipients with API ≥ 1.1, compared with median 2571 days (228-4661) in 9/39 (23%) recipients with API < 1.1 (p=0.008, K-M test) (Figure 2 and Figure S2). In the LT subcohort, graft loss occurred in 2/32 (6%) recipients with API ≥ 1.1 and 2/47 (4%) recipients with API < 1.1 (p=NS).

## Discussion

This study expands on our previous findings by showing that alloantigen uptake is representative of alloantigen presentation by B-cells and the downstream T-cell response (Figure 1a). The uptake of alloantigen was associated with enhanced Th activation resulting in higher frequencies of CD154+Th cells. The antigen presentation process encompasses antigen uptake and presentation (12-15). Inhibition of either process with cytochalasin and primaquine respectively, resulted in decreased frequencies of CD154+Th cells, compared with those seen in the absence of these inhibitors.

In our previous study, we found that an antigen presenting index (API), which measured the presentation of donor antigen relative to presentation of HLA-non-identical third-party or reference alloantigen distinguished rejectors from non-rejectors in a modest cohort of liver or intestine transplant recipients (11). Here, we further show using LR and ROC analysis, that a threshold API of 1.1 or greater can identify liver or intestine recipients who develop acute cellular rejection within the 60-day post-test period with a sensitivity, specificity, and positive and negative predictive values approaching or exceeding 80% in the validation set of post-transplant samples (Figures S1). For 20 pre-transplant samples, the prediction or rejection within the 60-day post-transplant period approaches or exceeds 75% for each performance metric. Applied to post-transplant samples for each organ system separately, an API of 1.1 or greater has PPV and NPV of 77% and 92%, respectively, for liver recipients, and 100% and 80% respectively, for intestine recipients (Figure S3 and Table S2). These findings have clinical application for the rapid assessment of rejection-risk. The test system and the reporting of the API can be completed in 4-6 hours. This can allow the caregiver to develop a treatment plan, while waiting for confirmatory biopsies, or validate the need for a confirmatory biopsy.

The added clinical benefits of measuring B-cell alloantigen presentation in the clinic may include identification of recipients at risk to develop DSA late after transplantation, and those who are at risk for late graft loss (Figure 2 and Figure S2). In our study, the API threshold of 1.1 or greater predicted a 2.2-fold greater risk of developing late DSA, one or more years after API measurements (Figure 2 and Figure S2). This threshold was also associated with a significantly higher incidence of late graft loss among intestine recipients, who are susceptible to this event because of the highly immunogenic intestine allograft. Liver transplant recipients did not demonstrate this association between B-cells that present donor and graft loss, likely because of the liver’s regenerative ability in the face of immunological injury. The added benefit of predicting who might develop late DSA can aid in the selection of maintenance regimens in those with an API ≥ 1,.and who may have experienced T-cell mediated coupled with circulating DSA. Such individuals are at risk for recurrent episodes of rejection. A high API may also stratify high-risk intestine transplant recipients into those who need more or less immunosuppression. After intestine transplantation, graft survival has yet to approach those of other organs.

We acknowledge the obvious limitations of our cross-sectional study, the modest sample size of a predominantly pediatric population, and the lack of serial measurements to describe a longitudinal relationship between changing immunosuppression levels in the course of routine care. Additional studies can address these limitations. In conclusion, donor antigen presentation by B-cells is associated with acute cellular rejection with clinically acceptable sensitivity and specificity, and may have the added benefit of predicting the development of DSA and late graft loss after liver or intestine transplantation.

## Acknowledgments

Techniques are described in patent US10222374, Assignee University of Pittsburgh, Licensee Plexision, Inc, Pittsburgh, PA, USA

Raizman-Hainey Endowed Fund, Hillman Foundation of Pittsburgh.

## Alphabetical List of All Abbreviations and Definitions

(Th): T-helper cells
(Tc): T-cytotoxic cells
(DSA): Donor-specific anti-HLA antibodies
(MHC): Major histocompatibility complex
(CFSE): Carboxyfluoresciensuccinimidylester
(PBL): Peripheral blood lymphocytes
(API): Antigen presenting index
(LR): Logistic regression
(ROC): Receiver-operating-characteristic
(FKWB): Tacrolimus whole blood concentration
(CI): Confidence intervals
(Cyto D): Cytochalasin D
(Prima): Primaquine
(AUC): Area under the receiver operating characteristic curve
(Tx): Transplantation
(PPV): Predictive Value
(NPV): Negative Predictive Value
(LT): Liver Transplantation
(IT): Intestine Transplantation

## Supplementary Figure Legends

**Figure S1: Test performance**. *Upper row*. Dot plots show API values for rejectors and non-rejectors in a) all post-transplant training set samples. These dot plots are shown in Figure 3 and are included here for reference, b) all post-transplant validation set samples and e) all pre-transplant validation set samples. *Lower row*. Area under the receiver operating characteristic curve (AUC) for corresponding APIs in each cohort.

**Figure S2: Association of DSA and graft loss with test result**. Test results are categorized as API>1.1 (red line) and API <1.1 (green line). *Upper row*. Time to late donor-specific anti-HLA antibody (DSA) measurements a. 85 of 131 total study subjects for whom DSA measurements were available (shown in Figure 3). b. Fifty five LT recipients (shown in Figure 3), and c. Thirty IT recipients. *Lower row*. Time to graft loss a. 131 total subjects, b. Seventy nine LT recipients, and c. Fifty two IT recipients (shown in Figure 3) (Kaplan-Meir test). Each K-M plot shows two subgroups, recipients with API <1.1 (green) and those with API ≥ 1.1 (red).

**Figure S3: Test performance by organ type**. *Upper row*. Dot plots show API values for rejectors and non-rejectors in a b) all liver transplant subjects from the training set, c) all intestine transplant recipients from the training set. *Lower row*. Area under the receiver operating characteristic curve (AUC) for corresponding APIs in each cohort.

## References

1. Marshall LS, Aruffo A, Ledbetter JA, et al. The molecular basis for T cell help in humoral immunity: CD40 and its ligand, gp39. J Clin Immunol 1993; 13: 165.

2. Gordon J, Katira A, Holder M, et al. Central role of CD40 and its ligand in B lymphocyte responses to T-dependent antigens. Cell Mol Biol (Noisy-le-grand) 1994; 40: 1.

3. M.J. Bevan. Helping the CD8+ T-cell response. Nat. Rev. Immunol., 4 (2004), pp. 595–602.

4. Grammer AC, Bergman MC, Miura Y, et al. The CD40 ligand expressed by human B cells costimulates B cell responses. J Immunol 1995; 154: 4996.

5. Grammer AC, McFarland RD, Heaney J, et al. Expression, regulation, and function of B cell-expressed CD154 in germinal centers. J Immunol 1999; 163: 4150.

6. S.P. Schoenberger, R.E.M. Toes, E.I.H. van der Voort, R. Offringa, C.J.M. MeliefT-cell help for cytotoxic T lymphocytes is mediated by CD40–CD40L interactions. Nature, 393 (1998), pp. 480–483.

7. S.R.M. Bennett, F.R. Carbone, F. Karamalis, R.A. Flavell, J.F.A.P. Miller, W.R. Heath. Help for cytotoxic-T-cell responses is mediated by CD40 signalling. Nature, 393 (1998), pp. 478–480.

8. Ashokkumar C, Bentlejewski C, Sun Q, Higgs BW, Snyder S, Mazariegos GV, Abu-Elmagd K, Zeevi A, Sindhi R. Allospecific CD154+ B cells associate with intestine allograft rejection in children. Transplantation 2010 Dec;90(11):1226–31. doi: 10.1097/TP.0b013e3181f995ce. PMID: 20881665.

9. Mellins ED, Stern LJ. HLA-DM and HLA-DO, key regulators of MHC-II processing and presentation. Curr Opin Immunol. 2014 Feb;26:115–22. doi: 10.1016/j.coi.2013.11.005. Epub 2013 Dec 8. Review. PMID: 24463216.

10. Yin L, Calvo-Calle JM, Dominguez-Amorocho O, Stern LJ. HLA-DM constrains epitope selection in the human CD4 T cell response to vaccinia virus by favoring the presentation of peptides with longer HLA-DM-mediated half-lives. J Immunol. 2012 Oct 15;189(8):3983–94. doi: 10.4049/jimmunol.1200626. Epub 2012 Sep 10.

11. Ningappa M, Ashokkumar C, Higgs BW, Sun Q, Jaffe R, Mazariegos G, Li D, Weeks DE, Subramaniam S, Ferrell R, Hakonarson H, Sindhi R. Enhanced B cell alloantigen presentation and its epigenetic dysregulation in liver transplant rejection. Am J Transplant. 2016 Feb;16(2):497–508. doi: 10.1111/ajt.13509. PMID: 26663361. PMCID: PMC5082419 [Available on 2017-02-01].

12. Telega GW, Baumgart DC, Carding SR. Uptake and presentation of antigen to T cells by primary colonic epithelial cells in normal and diseased states. Gastroenterology. 2000 Dec;119(6):1548–59.

13. Lanzavecchia A. Receptor-mediated antigen uptake and its effect on antigen presentation to class II-restricted T lymphocytes. Ann Rev Immunol 1990; 8: 773–793.

14. Levine TP, Chain BM. Endocytosis by antigen presenting cells: Dendritic cells are as endocytically active as other antigen presenting cells. Proc Natl Acad Sci U S A 1992; 89: 8342–8346.

15. Kakiuchi T, Chesnut RW, Grey HM. B cells as antigen-presenting cells: The requirement for B cell activation. J Immunol 1983; 31: 109–114.

16. Sindhi R, Higgs BW, Weeks DE, Ashokkumar C, Jaffe R, Kim C, Wilson P, Chien N, Glessner J, Talukdar A, Mazariegos G, Barmada MM, Frackleton E, Petro N, Eckert A, Hakonarson H, Ferrell R. Genetic variants in major histocompatibility complex-linked genes associate with pediatric liver transplant rejection. Gastroenterology. 2008 Sep;135(3):830-9, 839.e1-10. doi: 10.1053/j.gastro.2008.05.080. Epub 2008 Jun 3. PMID: 18639552; PMCID: PMC2956436.

17. Denzin LK, Sant’Angelo DB, Hammond C, Surman MJ, Cresswell P. Negative regulation by HLA-DO of MHC class II-restricted antigen processing. Science 1997 Oct;278(5335):106-09. PMID: 9311912.

18. Wozniak LJ, Hickey MJ, Venick RS, Vargas JH, Farmer DG, Busuttil RW, McDiarmid SV, Reed EF. Donor-specific HLA Antibodies Are Associated With Late Allograft Dysfunction After Pediatric Liver Transplantation. Transplantation. 2015 Jul;99(7):1416–22. doi: 10.1097/TP.0000000000000796. PMID: 26038872; PMCID: PMC5283576.

19. Kaneku H, O’Leary JG, Banuelos N, Jennings LW, Susskind BM, Klintmalm GB, Terasaki PI. De novo donor-specific HLA antibodies decrease patient and graft survival in liver transplant recipients. Am J Transplant. 2013 Jun;13(6):1541–8. doi: 10.1002/ajt.12212. PMID: 23721554; PMCID: PMC4408873.

20. Ashokkumar C, Green M, Soltys K, Michaels M, Mazariegos G, Reyes-Mugica M, Higgs BW, Spishock B, Zaccagnini M, Sethi P, Rzempoluch A, Kepler A, Kachmar P, Remaley L, Winnier J, Jones K, Moir K, Fazzolare T, Jenkins K, Hartle T, Falik R, Ningappa M, Bond G, Khanna A, Ganoza A, Sun Q, Sindhi R. CD154-expressing CMV-specific T cells associate with freedom from DNAemia and may be protective in seronegative recipients after liver or intestine transplantation. Pediatr Transplant. 2020 Feb;24(1):e13601. doi: 10.1111/petr.13601. Epub 2019 Oct 27. PMID: 31657119.

21. Ashokkumar C, Soltys K, Mazariegos G, Bond G, Higgs BW, Ningappa M, Sun Q, Brown A, White J, Levy S, Fazzolare T, Remaley L, Dirling K, Harris P, Hartle T, Kachmar P, Nicely M, O’Toole L, Boehm B, Jativa N, Stanley P, Jaffe R, Ranganathan S, Zeevi A, Sindhi R. Predicting Cellular Rejection With a Cell-Based Assay: Preclinical Evaluation in Children. Transplantation. 2017 Jan;101(1):131–140. doi: 10.1097/TP.0000000000001076. PMID: 26950712; PMCID: PMC5011025.

22. Ashokkumar C, Talukdar A, Sun Q, Higgs BW, Janosky J, Wilson P, Mazariegos G, Jaffe R, Demetris A, Dobberstein J, Soltys K, Bond G, Thomson AW, Zeevi A, Sindhi R. Allospecific CD154+ T cells associate with rejection risk after pediatric liver transplantation. Am J Transplant. 2009 Jan;9(1):179–91. doi: 10.1111/j.1600-6143.2008.02459.x. Epub 2008 Oct 31. PMID: 18976293; PMCID: PMC2997472.

