## Supplementary Tables for "Enhanced donor antigen presentation by B-cells predicts acute cellular rejection and late outcomes after transplantation"

**Table S1.** Characteristics of training and validation set subjects. All=all subjects and samples tested. Tx=transplantation. $API_0$=pre-transplant sample. $API_1$= sample obtained in the first 60 days post-transplantation. $API_x$= sample 61 days or more post-transplantation **(We should add the type of organ liver, intestine, Liver intestine or Liver kidney).**

| | Test set | Validation set | p- value | $API_0$ |
|---|---|---|---|---|
| | $API_1$ *and* $API_x$ | $API_1$ *and* $API_x$ | test vs validation $API_1$ *and* $API_x$ | Unique Subjects |
| **Parameters** | **All** | **All** | | |
| Total subjects | 99 | 32 | | 7 |
| Age at Tx (mean±SD) | 6.02 ± 6.32 | 6.86 ± 6.08 | 0.511 (NS) | 6.74 ± 8.2 |
| Age (Range) years | 0.01 to 23.94 | 0.06 to 19.09 | | 0.6 to 23.96 |
| Gender (M:F) | 54:45 | 19:13 | 0.686 (NS) | 4:3 |
| Race (Caucasian vs. non-Caucasian) | 82:17 | 25:7 | 0.601 (NS) | 7:0 |
| Organ: Liver : Intestine | 54:45 | 25:7 | 0.067 (NS) | 5:2 |
| Induction: No-induction : Induction | 32:67 | 16:16 | 0.092 (NS) | 5:2 |
| FKWB ( ng / ml ) | 8.46 ± 7.20 | 6.07 ± 2.94 | 0.011 | NA |
| $API_1$ samples | 29 | 7 | 0.499 (NS) | NA |
| $API_x$ samples | 70 | 25 | | NA |
| Time between Tx and sample (days) $API_1$ and $API_x$ | 1063 ± 1479 | 1964 ± 1907 | 0.019 | -0.6 ± 1.3 |
| (range) | (5, 5817) | (14, 5567) | | (-3, 1) |
| Time between sample and biopsy/event (days) | 6.65 ± 12.17 | 3.00 ± 7.42 | 0.237 (NS) | 31.3 ± 14.5 |
| (range) | (0, 50) | (0, 30) | | (15, 59) |

**Table S2.** Test performance based on organ type in training set samples, and in the validation sets of post-transplant and pre transplant samples, based on the API threshold of 1.1 or greater. Unless otherwise stated, post-transplant samples were used in the training set and validation set. PPV-positive predictive value, NPV-negative predictive value, CI-confidence interval

| Training set | LTx (n=54) | 95% CI | ITx (n=45) | 95% CI |
|---|---|---|---|---|
| Sensitivity | 0.9 (18/20) | 0.67, 0.98 | 1.00 (10/10) | 0.67, 1.00 |
| Specificity | 0.97 (33/34) | 0.83, 1.00 | 0.97 (34/35) | 0.84, 1.00 |
| PPV | 0.95 (18/19) | 0.72, 1.00 | 0.91 (10/11) | 0.60, 1.00 |
| NPV | 0.94 (33/35) | 0.80, 1.00 | 1.00 (34/34) | 0.88, 1.00 |

| Validation set | LTx (n=25) | 95% CI | ITx (n=7) | 95% CI |
|---|---|---|---|---|
| Sensitivity | 0.91 (10/11) | 0.57, 1.00 | 0.67 (2/3) | 0.13, 0.98 |
| Specificity | 0.79 (11/14) | 0.50, 0.95 | 1.00 (4/4) | 0.40, 1.00 |
| PPV | 0.77 (10/130 | 0.46, 0.94 | 1.00 (2/2) | 0.2, 1.00 |
| NPV | 0.92 (11/12) | 0.60, 1.00 | 0.80 (4/5) | 0.3, 0.99 |

| Validation set (pre-transplant) | LTx (n=14) | 95% CI | ITx (n=06) | 95% CI |
|---|---|---|---|---|
| Sensitivity | 0.75 (6/8) | 0.36, 0.96 | 0.67 (2/3) | 0.13, 0.98 |
| Specificity | 1.00 (6/6) | 0.52, 1.00 | 1.00 (3/3) | 0.31, 1.00 |
| PPV | 1.00 (6/6) | 0.52, 1.00 | 1.00 (2/2) | 0.20, 1.00 |
| NPV | 0.75 (6/8) | 0.36, 0.96 | 0.75 (3/4) | 0.22, 0.99 |
