## Supplementary Figures for "Enhanced donor antigen presentation by B-cells predicts acute cellular rejection and late outcomes after transplantation"

### Slide 1
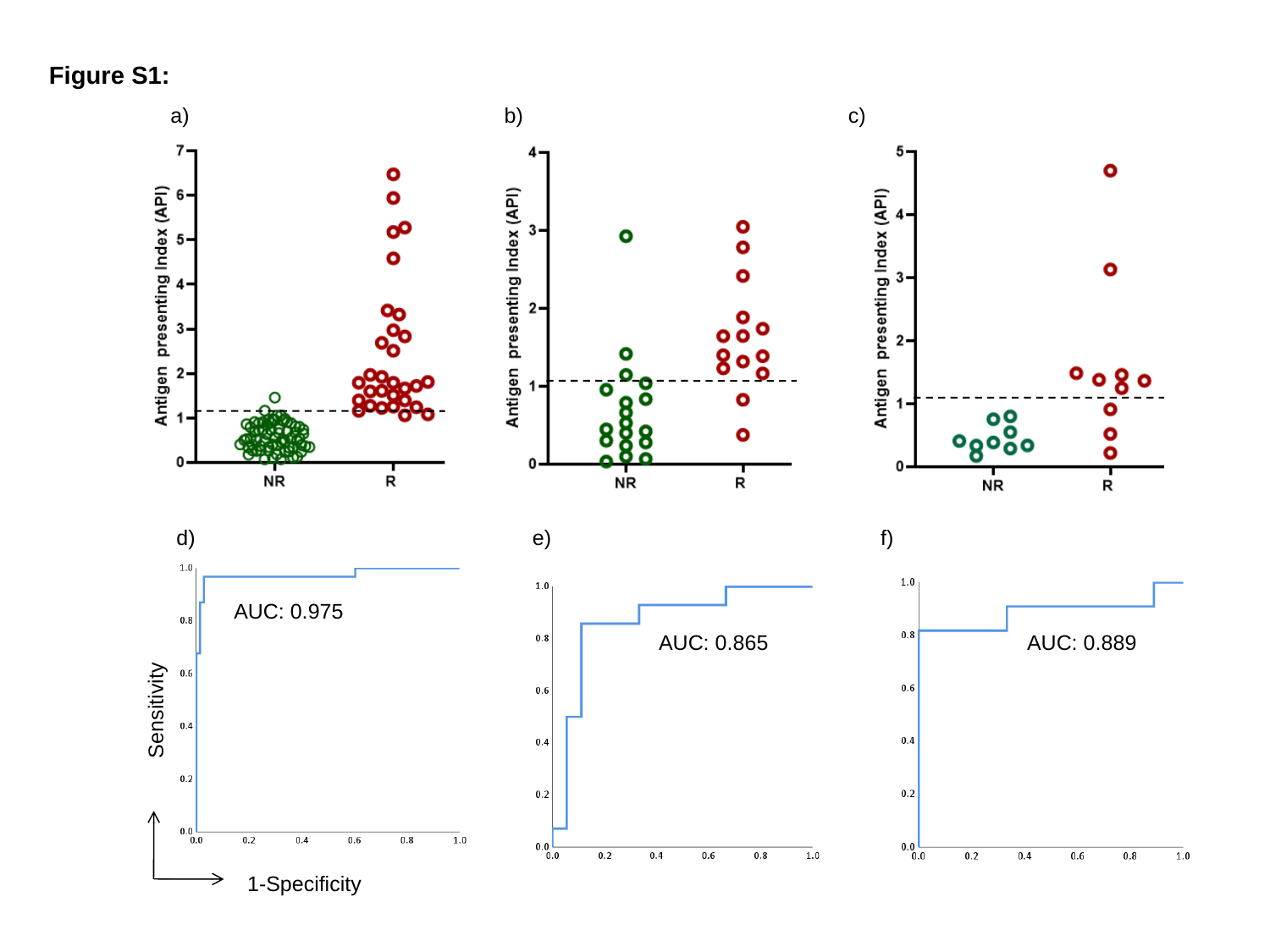

Figure S1:
a)
b)
c)
d)
e)
f)
AUC: 0.975
AUC: 0.865
AUC: 0.889
Sensitivity
1-Specificity

### Slide 2
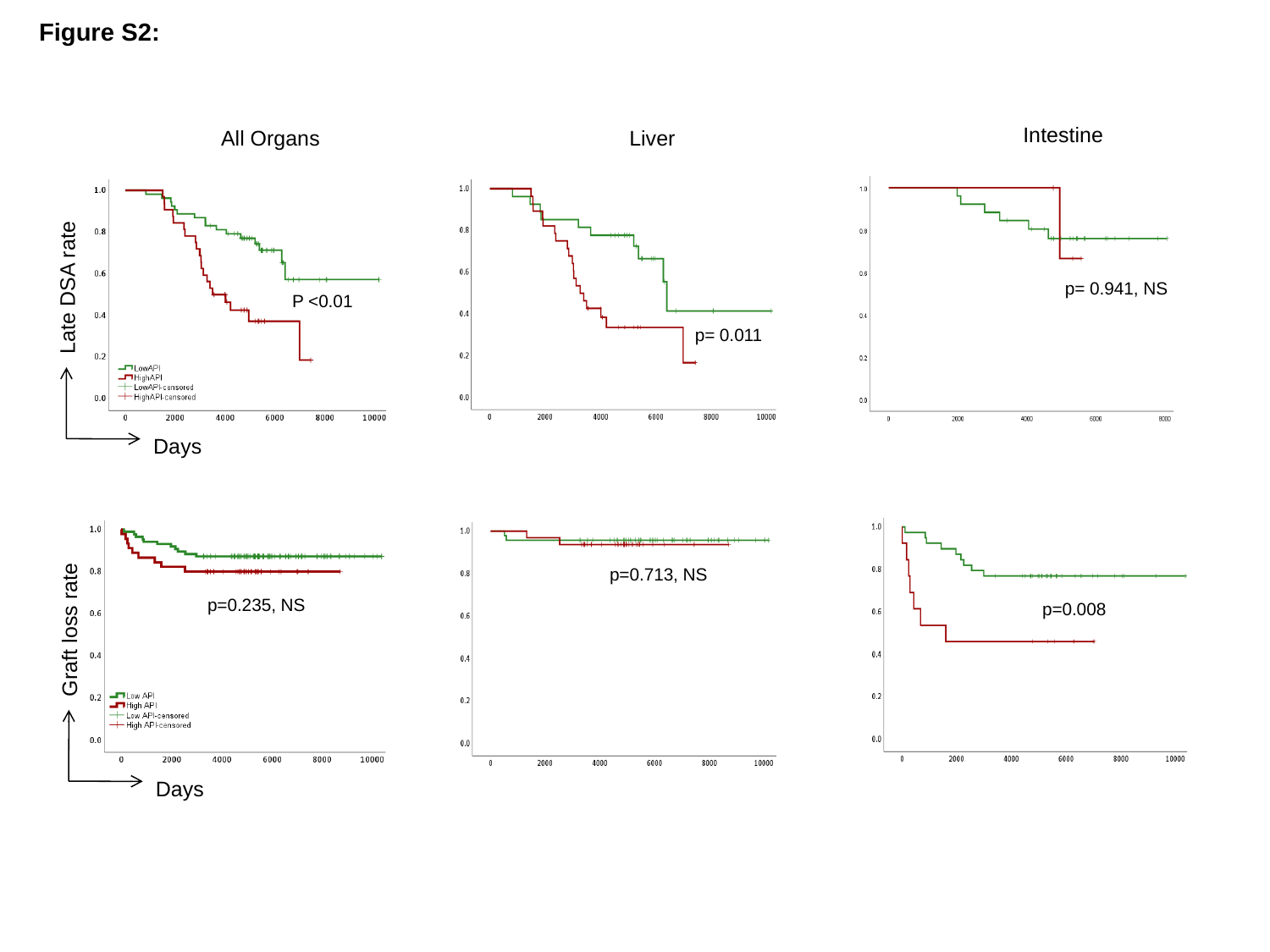

Figure S2:
Intestine
All Organs
Liver
Late DSA rate
p= 0.941, NS
P <0.01
p= 0.011
Days
p=0.713, NS
p=0.235, NS
p=0.008
Graft loss rate
Days

### Slide 3
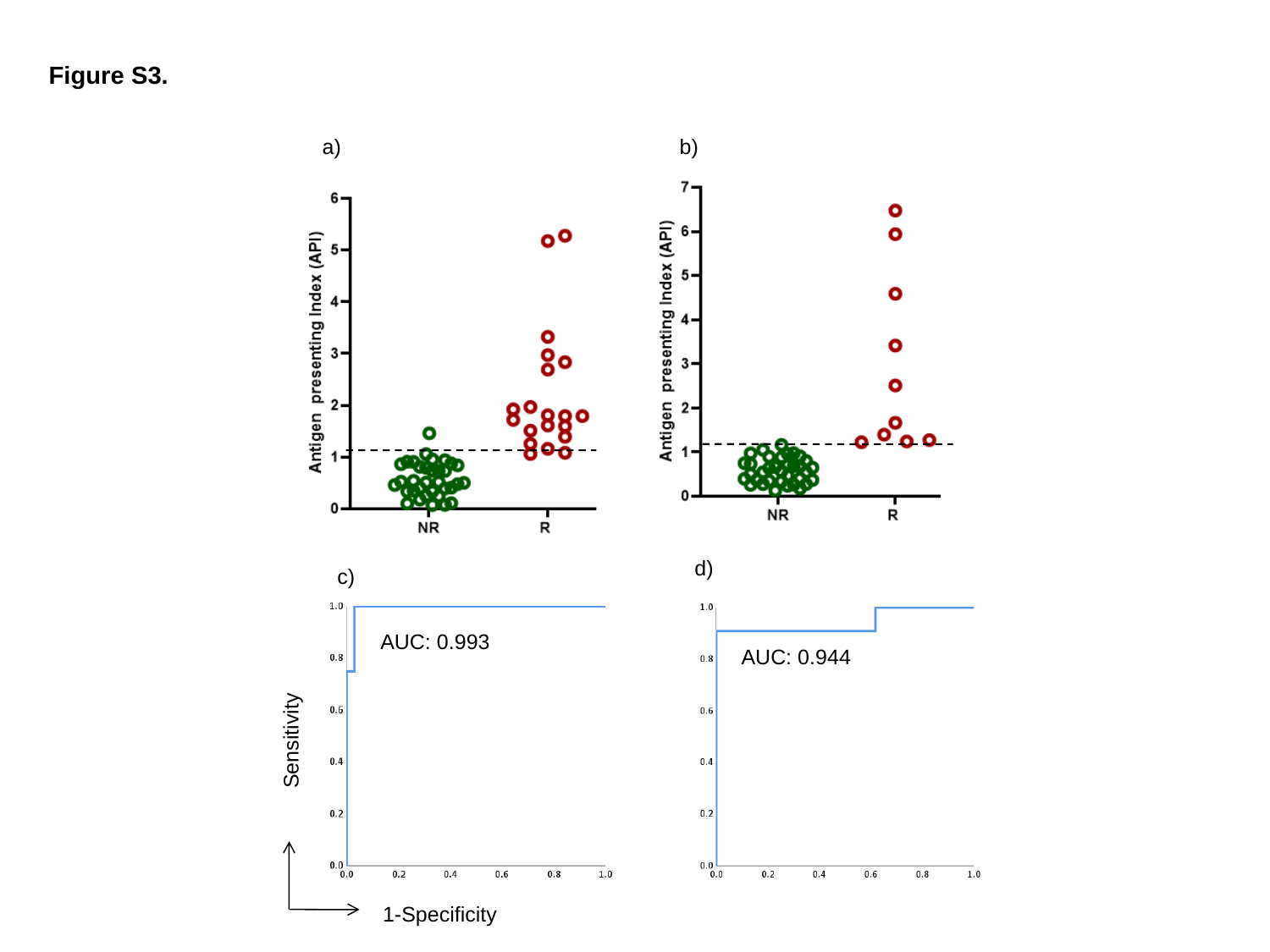

Figure S3.
a)
b)
d)
c)
AUC: 0.993
AUC: 0.944
Sensitivity
1-Specificity
